# Benchmarking water-limited yield potential and yield gaps of Shiraz in the Barossa and Eden Valleys

**DOI:** 10.1101/2022.10.25.513680

**Authors:** Marcos Bonada, Vinod Phogat, Cassandra Collins, Paul R. Petrie, Victor Sadras

## Abstract

**Background and Aims:** Vineyard performance is impacted by water availability including the amount and seasonality of rainfall and evapotranspiration and irrigation volume. We benchmarked water-limited yield potential (Yw), calculated yield gaps as the difference between Yw and actual yield, and explored the underlying environmental and management causes of these gaps.

**Methods and Results:** The yield and its components in two sections of 24 Shiraz vineyards was monitored during three vintages in the Barossa zone (GI). The frequency distribution of yield was L-shaped, with half the vineyards below 5.2 t ha^-1^, and an extended tail of the distribution that reached 24.9 t ha^-1^. The seasonal ratio of actual crop evapotranspiration and reference evapotranspiration was below 0.48 in 85% of cases, with a maximum of 0.65, highlighting a substantial water deficit in these vineyards. A boundary function relating actual yield and seasonal rainfall was fitted to quantify Yw. Yield gaps increased with increasing vine water deficit, quantified by the carbon isotope composition in the fruit. The yield gap was smaller with higher rainfall before budburst, putatively favouring early-season vegetative growth and allocation to reproduction, and with higher rainfall between flowering and veraison, putatively favouring fruit set and berry growth. The gap was larger with higher rainfall and lower radiation between budburst and flowering. The yield gap increased linearly with vine age between 6 and 33 yr at a rate of 0.3 t ha^-1^ yr^-1^. The correlation between yield gap and yield components ranked bunch weight ≈ berries per bunch > bunch number > berry weight; the minimum to close the yield gap was 185,000 bunches ha^-1^, 105 g bunch^-1^, 108 berries bunch^-1^ and 1.1 g berry^-1^.

**Conclusions:** Water deficit and vine age were major causes of yield gaps. Winter irrigation provides an opportunity to improve productivity. The cost of dealing with older, less productive vines needs to be weighed against the rate of increase in yield gap with vine age.

**Significance of the Study:** A boundary function to estimate water-limited yield potential returned viticulturally meaningful yield gaps and highlighted potential targets to improve vineyard productivity.

## Introduction

Mediterranean regions feature at least 60% of annual rainfall in the winter half-year (di Castri and Mooney 1973). The Barossa and Eden Valley regions, the focus of this study, have a Mediterranean-type climate where vine growth and yield rely on three sources of water: soil-stored winter rainfall, variable amounts of summer rainfall from the tails of tropical storms, and supplementary irrigation (Dry, et al. 2005). Climate change is placing increased stress on water resources and supports a revisions of water management strategies and how they relate to vineyard productivity. First and most important, winter rainfall is diminishing in South-eastern Australia (Cai and Cowan 2013), and irrigation approaches are required to account for drier soil in spring (Bonada, et al. 2021, Bonada, et al. 2020). Second, irrigation is also being increasingly used to promote evaporative cooling as the central component to the management of heat waves, which are increasing in frequency and intensity (Sadras and Schultz 2012, Webb, et al. 2010). Third, decompressing harvest by using pruning after budburst is being used to displace critical developmental stages onto cooler conditions, and requires further adjustments in irrigation to account for shifts in the dynamics of canopy cover (Moran, et al. 2019, Moran, et al. 2018, Moran, et al. 2021, Petrie, et al. 2017). For example, the evapotranspiration of Malbec in Mendoza was 5% lower in vines pruned at 2-3 separated leaves, and 10% lower in vines pruned at 8 separated leaves than in winter-pruned controls (Morgani, et al. 2022).

Production functions relating crop yield and water use (or irrigation application) are critical for irrigation management (Steduto, et al. 2012). These functions are usually tight, i.e., high r^2^, because yield and water use are measured in experiments where water supply is the main source of variation and other factors are largely controlled (Steduto, et al. 2012). In contrast, the association between yield and water use is scattered where multiple, agronomically relevant sources of variation influence crop traits; for rainfed wheat in Australia, for example, the relationship between yield and water use has a typical r^2^ ∼ 0.3 (Sadras 2020). Dealing with a highly scattered yield-water use relation, French and Schultz (1984) insightfully fitted a boundary function with biophysically meaningful parameters, rather than a regression. Their boundary function is

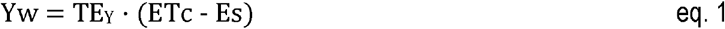

where Yw is water-limited yield potential, ETc is actual crop evapotranspiration, the slope TE_Y_ is the maximum transpiration efficiency for yield, and the x-intercept Es is an approximate measure of non-productive water loss, primarily soil evaporation. This model has many simplifications, including a constant soil evaporation and lack of consideration of the effect of timing of water supply on yield (French and Schultz 1984, Rockström 2003, Sadras, et al. 2015).

In a similarly simple, one equation model, Rockström (2003) relates ETc per unit yield and yield, hence accounting for the decline in Es:ETc with increasing crop vigour and yield:

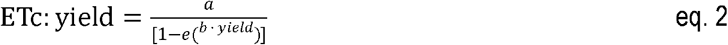

where *a* is the transpiration-to-yield ratio and *b* is the rate of decline in soil evaporation (Es) with increased canopy size, and therefore also the yield at which Es:ETc reaches its minimum. This model has an element of circularity because yield is common to *x* and *y* (Brett 2004). In grapevine, both these models (eq. 1 and eq. 2) further overlook yield response to water supply in the previous season, but so do the typical production functions that consider water use in the current season only (Steduto, et al. 2012) and more refined models of grapevine growth and yield (Yang, et al. 2022 and references cited therein).

Owing to its transparency and frugal data requirement, eq. (1) has been widely adopted in the industry where cereal farmers relate their actual yield (Ya) to the water-limited yield potential, calculate a yield gap, i.e., Yw – Ya, identify its causes, and modify their practices to close the gap (Sadras 2020). The yield gap, its causes, and remedies have motivated a global yield gap initiative that accounts for data-rich and data-poor cropping systems (https://www.yieldgap.org/). Daily time-step models accounting for crop, soil, climate, and management are favoured to estimate the water-limited yield potential, hence overcoming the limitations of simpler approaches (van Ittersum, et al. 2013). Owing to carry over effects across seasons affecting processes such as bud fertility that in turn drive bunch number and the dynamics of carbohydrate reserves buffering berry growth (Dunn, et al. 2004, Iland, et al. 2011, Vasconcelos, et al. 2009), reliable models to predict yield are lagging and yield gap analysis is incipient in grapevine. For example, Yang et al. (2022) modelled a single yield component, bunch weight, to calculate yield gaps in European wine regions under the explicit assumption of no variation in both plant population density and number of bunches per ha.

In this study, we benchmark the water-limited yield potential, and calculate yield gaps of Shiraz in the Barossa regions using a data set of actual yield that captures viticulturally relevant sources of variation including weather, soil, vineyard age, and management.

## Methods

### Study area and experimental sites

This study was conducted in the Barossa Zone GI, which includes the Barossa and Eden Valley regions. The zone is internationally known for its full-bodied Shiraz wines. The complex system of valleys and twisting hills results in a variety of slopes, aspects and sites (Bramley and Ouzman 2021). The soils vary widely due to the complex geology of the region, but they primarily fall in a family of duplex soils, characterised by an abrupt texture change between the topsoil and subsoil (Dry, et al. 2005).

We measured yield and its components, and calculated actual crop evapotranspiration in 144 samples that resulted from the full combination of six sub-regions, four vineyards per subregion, two sections per vineyard, and three consecutive vintages since 2019. The subregions are Northern Grounds (NG), Central Grounds (CG), Eastern Edge (EE), Southern Grounds (SG), Western Ridge (WR) and Eden Valley (EV) (www.barossawine.com/vineyards/barossa-grounds/). The vineyard sites were divided into two sections (high, low) based on an electromagnetic induction soil survey and canopy size maps of each block, as explained in Bonada et al. (2022). Vineyard age, clone, rootstock, row orientation, pruning method, and trellising system were recorded (Supplementary Table 1).

### Vine traits

At harvest, the number of bunches and yield per metre of cordon were recorded from three sections within each sample location (high, low). We calculated average bunch weight from yield and bunch number and the number of berries per bunch from bunch weight and average berry weigh from a 100-berry sample. In winter, we counted the number of shoots and measured pruning weight in three one-meter canopy sections. Yield, bunch number, shoot number and pruning weight per ha were calculated based on the vine and row spacing (Supplementary Table 1).

To quantify crop water status, we measured carbon isotope composition (δ^13^C) in must. This trait integrates crop water status over the growing period until sampling time and is robust in relation to environmental conditions—radiation, wind speed, temperature, vapour pressure deficit (Condon, et al. 2002) – unlike traits such as stomatal conductance, leaf water potential, or canopy temperature that vary with conditions at sampling time. We measured δ^13^C in 144 samples of must. Juice collected at harvest was centrifuged for 4 minutes at 4500 RPM and 50 ml were sterilised the same day by autoclaving at 120 °C for 20 minutes (Ollat, et al. 2002). Carbon isotope composition was measured on approximately 10 μl of berry juice, freeze dried in tin capsules (3 mg dried weight) using a continuous flow isotope ratio mass spectrometer (Nu Horizon IRMS with EuroVector EA, Wrexham, United Kingdom).

### Weather, soil moisture and water balance

Bonada et al. (2022) describe the measurements of weather and components of the soil water balance in detail. Briefly, 24 weather stations (MEA Junior WS, Magill, South Australia), one at each site, were installed at the beginning of the experiment to log temperature, relative humidity, wind speed and direction, solar radiation, and rainfall at 15-minutes intervals; daily reference evapotranspiration (ETo) was calculated with the method of Penman–Monteith {Allen, 1998 #4188}. Soil moisture was continually monitored at four depths using capacitance probes (EnviroSCAN system, Sentek, Magill, South Australia). The PVC access tubes were installed directly below the irrigation lateral and between drippers. Data were logged every 15 minutes and retrieved from the logger at least monthly for processing using the IrriMax10 software (Sentek, Magill, South Australia). Details of probe calibration and calculations of plant available water are in Bonada et al. (2022). Crop evapotranspiration (ETc) and its components (transpiration, Tp; and evaporation, Es) were estimated using the FAO-56 dual-crop coefficient approach (Allen, et al. 1998) as described in Phogat et al. (2020).

### Statistical analyses

The analysis was constrained to 127 out of 144 paired yield-ET points due to missing samples of yield or ET components. We used an ANOVA to test the effect of main sources of variation on yield and its components, and other related traits. To explore associations between variables, we fitted least squares regression (LS, Model I) when error in x was negligible in comparison to error in y and reduced maximum axis regression (RMA, Model II) to account for error in both x and y (Niklas, 1994). Pearson’s correlation coefficients (r) were calculated, with *p* derived from Fisher z-transformation that transforms the sampling distribution of r for it to become normally distributed. For both ANOVA and regressions, we follow updated statistical recommendations and avoid the wording “statistically significant”, “non-significant”, or the variations thereof, thus avoiding dichotomisation based on an arbitrary discrete p-value (Wasserstein et al., 2019). Instead, we report p as a continuous quantity, and Shannon information transform [*s* = -log_2_(*p*)] as a measure of the information against the tested hypothesis (Greenland, 2019). Although *s* is a function of *p*, the additional information is not redundant. With the base-2 log, the units for measuring this information are bits (binary digits). For example, the chance of seeing all heads in 4 tosses of a fair coin is 1/24 = 0.0625. Thus, *p* = 0.05 conveys only *s* = -log_2_(0.05) = 4.3 bits of information, “which is hardly more surprising than seeing all heads in 4 fair tosses” (Greenland, 2019).

Boundary functions were fitted using percentile regression to define water-limited yield potential, and yield gaps were calculated as the difference between water-limited yield potential and actual yield as outlined in Sadras et al. (2015). To explore the associations of yield gap and weather, location, vineyard features, and management we used an ANOVA for discrete variables (e.g., trellising system), and regression for continuous variables (e.g., vine age).

## Results

### Growing conditions

Weather during the growing season varied with location, vintage, and intra-seasonally from budburst in spring to maturity in autumn (Fig. 1). Minimum temperature varied 4.9-fold, vapour pressure deficit varied 4.3-fold, and maximum temperature and solar radiation varied 2.1-fold.

**Fig 1.**
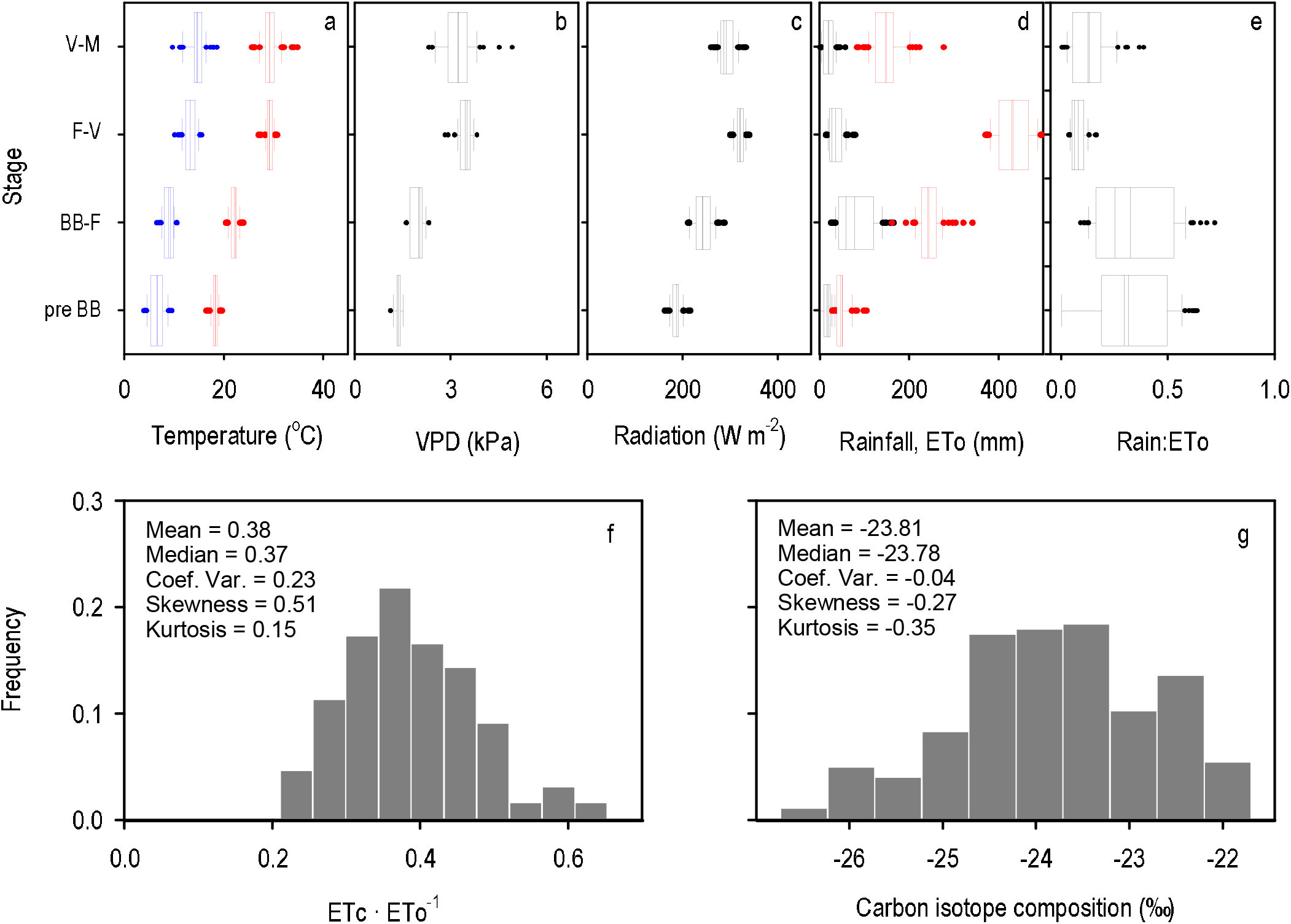
Frequency distribution of (a) minimum (blue) and maximum (red) temperature, (b) vapour pressure deficit, (c) solar radiation, (d) rainfall (black) and reference evapotranspiration (red), and (e) rain-to-reference evapotranspiration ratio during the periods from 29 August to budburst (pre-BB), from budburst to flowering (BB-F), from flowering to veraison (F-V), and from veraison to maturity (V-M). Frequency distribution of (f) the seasonal ratio between actual and reference evapotranspiration (ETc · ETo^-1^) and (g) carbon isotope composition in must, a trait related to vine water status.

Plant available water in the soil at budburst ranged from 27 to 144 mm and averaged 91 mm with a coefficient of variation of 0.26. Rainfall was below reference evapotranspiration, particularly between flowering and veraison (Fig. 1 d,e). Seasonal irrigation varied from 0 to 335 mm, and averaged 161 mm, with a coefficient of variation of 0.38. The seasonal ratio of actual crop evapotranspiration (accounting for plant available water in the soil at budburst, rainfall, and irrigation), and reference evapotranspiration was below 0.48 in 85% of cases, with a maximum of 0.65 (Fig. 1f), highlighting the prevalence of a substantial water deficit in these vineyards. Carbon isotope composition, a trait that relates to plant water status, varied from -26.74 to -21.70‰ (Fig. 1g).

### Yield and its components

Yield averaged 3.7 ± 0.37 t ha^-1^ in 2020, 5.9 ± 0.50 t ha^-1^ in 2019, and 10.1 ± 0.37 t ha^-1^ in 2021. For the pooled data, the frequency distribution of yield was L-shaped (Fig. 2a), with skewness and kurtosis departing from the normal distribution’s zero and three, respectively (Bradley 1982). Half the vineyards were below 5.2 t ha^-1^, and the extended tail of the distribution reached 24.9 t ha^-1^ (Fig. 2). Yield correlated with all its components (Table 1); the strength of the correlation declined in the order bunch number per ha, bunch weight, berries per bunch, berry weight. Yield correlated with vegetative traits including pruning weight and shoot number (Table 1). Yield, its components, and vegetative traits all declined with lower carbon isotope composition indicative of more severe water stress (Fig. 2b, Table 1).

**Table 1.** Correlation matrix of yield, its components, vegetative traits and carbon isotope composition in must (δ^13^C) for Shiraz in the Barossa zone (GI). Sources of variation are 25 vineyard locations, two sites per vineyard, and three vintages. Numbers in italics are Pearson’s correlation coefficients and numbers with coloured background are *p* from Fisher’s r to z test. *p* scale

|  |  |  |  |  |  |  |  |  |  |  |
| --- | --- | --- | --- | --- | --- | --- | --- | --- | --- | --- |
| 0.0001 | 0.001 | 0.01 | 0.05 | 0.1 | 0.2 | 0.3 | 0.4 | 0.5 | 0.6 | 1 |

| | Bunch no/ha | Bunch weight (g) | Berries/bunch | Berry weight (g) | Shoot no/m | Pruning wt (t/ha) | $\delta^{13}\text{C}(\text{‰})$ |
| --- | --- | --- | --- | --- | --- | --- | --- |
| Yield (t/ha) | *0.75* | *0.70* | *0.64* | *0.42* | *0.37* | *0.61* | *-0.36* |
| Bunch no/ha |  | *0.13* | *0.14* | *0.09* | *0.65* | *0.42* | *-0.30* |
| Bunch weight (g) |  |  | *0.91* | *0.55* | *-0.14* | *0.50* | *-0.28* |
| Berries/bunch |  |  |  | *0.20* | *-0.15* | *0.45* | *-0.19* |
| Berry weight (g) |  |  |  |  | *0.02* | *0.38* | *-0.26* |
| Shoot no/m |  |  |  |  |  | *0.43* | *-0.26* |
|  |  |  |  |  |  |  | *-0.18* |
| Yield (t/ha) | 0.0001 | 0.0001 | 0.0001 | 0.0001 | 0.0001 | 0.0001 | 0.0001 |
| Bunch no/ha |  | 0.0698 | 0.0542 | 0.2296 | 0.0001 | 0.0001 | 0.0001 |
| Bunch weight (g) |  |  | 0.0001 | 0.0001 | 0.0516 | 0.0001 | 0.0001 |
| Berries/bunch |  |  |  | 0.0054 | 0.032 | 0.0001 | 0.0079 |
| Berry weight (g) |  |  |  |  | 0.7929 | 0.0001 | 0.0002 |
| Shoot no/m |  |  |  |  |  | 0.0001 | 0.0116 |
| Pruning wt (t/ha) |  |  |  |  |  |  | 0.0002 |

**Fig 2.**
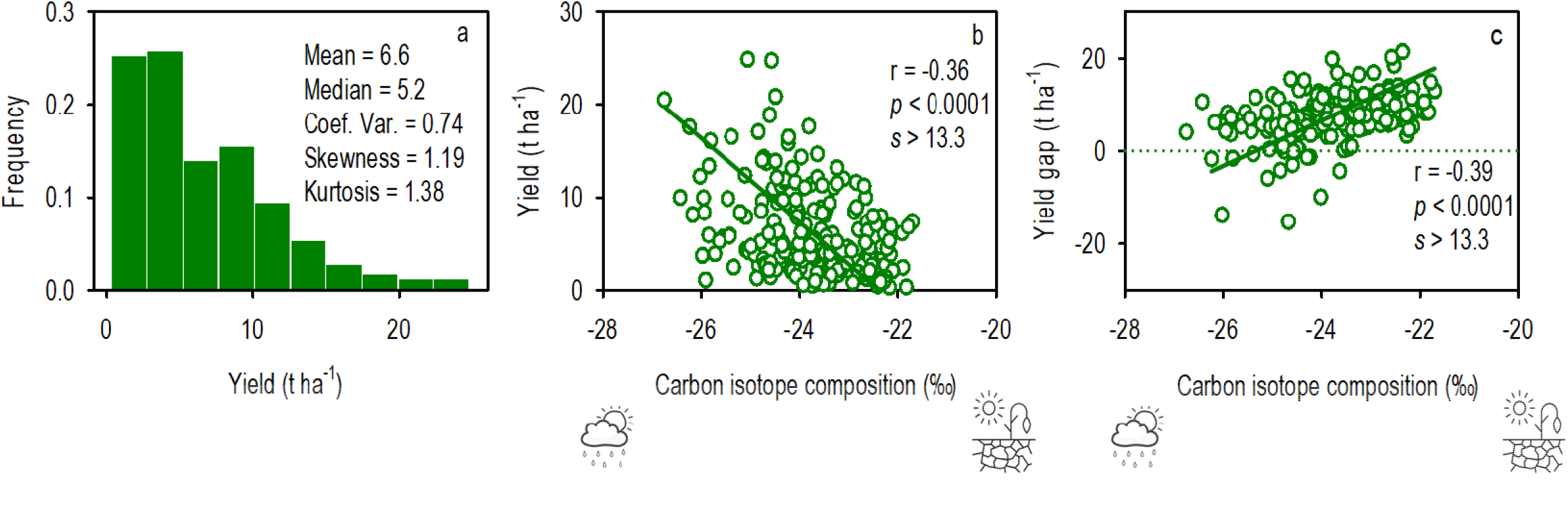
(a) Frequency distribution of yield of Shiraz in the Barossa zone (GI). (b) Relation between yield and carbon isotope composition. (c) Relation between yield gap and carbon isotope composition. In (b, c) lines are reduced maximum axis regressions.

### Relationship between ETc:yield ratio and yield

Rockström (2003) described the relation between the ratio ETc:yield and yield with eq. 2. He interpreted the declining ETc:yield ratio with increasing yield in terms of shifts in the evapotranspiration components: with increasing yield and crop vigour, the Es:ETc ratio declines, and a higher proportion of ETc is used in plant transpiration. The actual relation between ETc:yield and yield conformed to the expected model (Fig. 3a, r = 0.98, p < 0.0001, s > 13.3). However, this association involves a shared factor in y and x, with the potential for spurious correlations (Brett 2004). We tested the legitimacy of the association in Fig. 3a with two complementary approaches. Statistically, a large spurious correlation emerges when the coefficient of variation of the shared factor is more than 1.5 times larger than the coefficient of variation of the non-shared factor (Brett 2004). In our data set, the coefficient of variation of the shared factor, yield, was 73.8%, and the coefficient of variation of the non-shared factor, seasonal ETc, was 22.5%; the ratio is 3.3, indicating the spurious nature of the correlation from a statistical viewpoint. Biophysically, the association between Es:ETc and yield was weak and had a flat slope of 0.018 ± 0.002 (t ha^-1^)^-1^ (Fig. 3b). This analysis therefore supports the assumption of a conserved Es for the range of yield in our data and justifies a single relationship between yield and ETc (or rainfall) as summarised in eq. 1.

**Fig 3.**
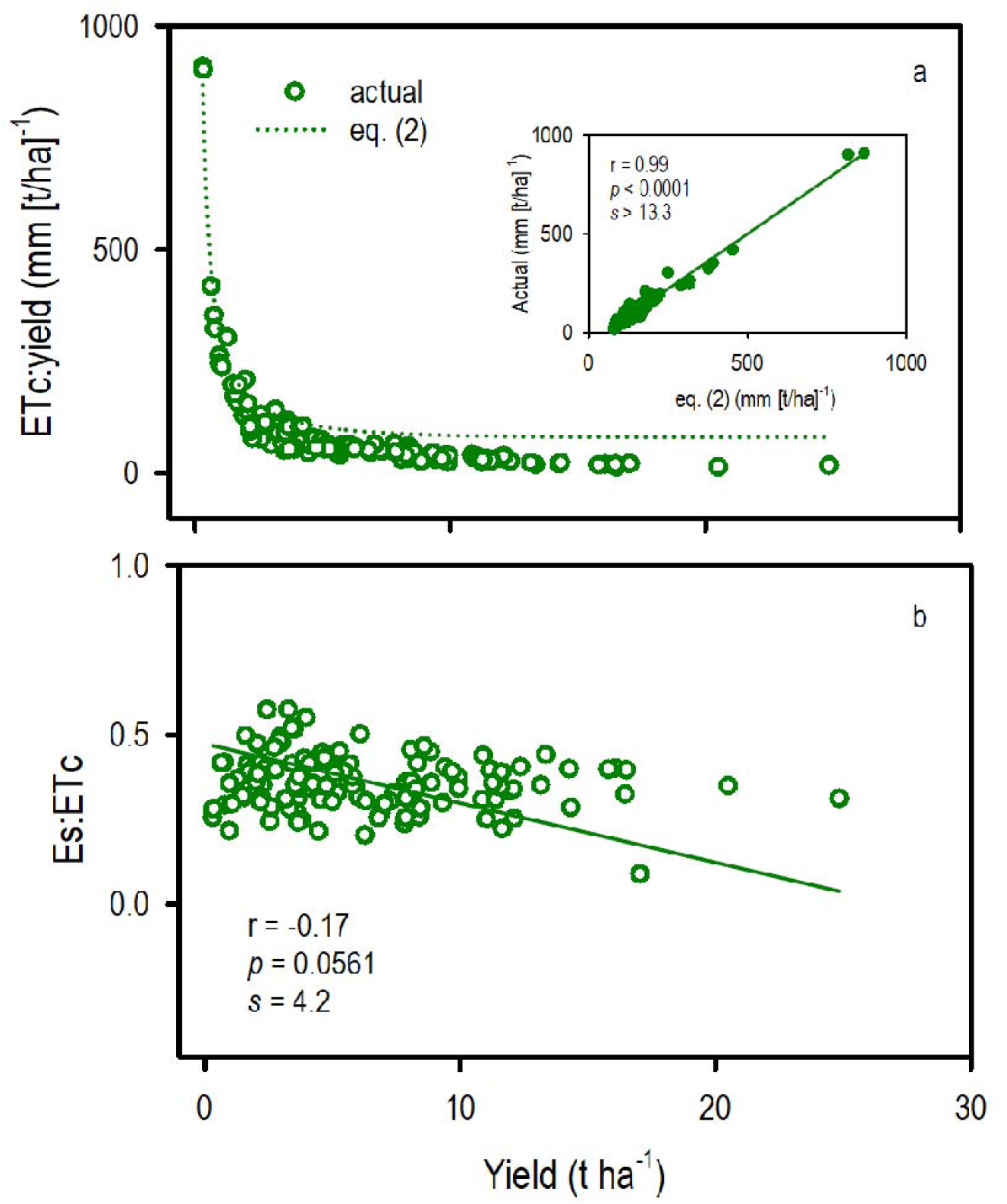
(a) Comparison of the actual relation between the ratio of crop evapotranspiration and yield and the model in eq. (2). The parameters for eq. (2) are *a* = 80 mm/t ha^-1^, *b* = -0.3 (Rockström 2003). In the inset, the line is the least-squares regression. (b) Relation between the ratio of soil evaporation and crop evapotranspiration and yield; the line is the reduced maximum axis regression. ETc: crop evapotranspiration, Es: soil evaporation.

### Benchmarking water-limited yield potential

The RMA regression between yield and seasonal ETc returned a slope representing yield per unit transpiration of 0.065 t ha^-1^ mm^-1^ and an x-intercept representing soil evaporation of 216 mm (Fig. 4a, Table 2). The strength of the relationship improved with ETc corrected by VPD, but not with ETc normalised with ETo (Fig. 4abc, Table 2). The RMA regression between yield and seasonal rainfall returned a slope of 0.117 t ha^-1^ mm^-1^ and an x-intercept of 74 mm (Fig. 4d, Table 2); this association (r = 0.62, F_1,124_ = 76.5) did not improve with corrections by VPD or ETo (Table 2), or with the inclusion of winter rainfall (r = 0.34, F_1,124_ = 16.7).

**Table 2.** Statistics of reduced maximum axis regressions between yield and ETc, ETc · VPD^-1^, ETc · ETo^-1^, rain, rain VPD^-1^ rain ETo-1^-1^ Scatterplots and fitted regressions are in Figure 4.

| Variable | r | F<sub>1,124</sub> | p | s | slope | x-intercept |
| --- | --- | --- | --- | --- | --- | --- |
| ETc (mm) | 0.32 | 13.99 | 0.0003 | 11.7 | 0.065 t ha<sup>-1</sup> mm<sup>-1</sup> | 216 mm |
| ETc · VPD<sup>-1</sup> (mm kPa<sup>-1</sup>) | 0.38 | 21.5 | 0.0001 | 13.3 | 0.160 t ha<sup>-1</sup> mm<sup>-1</sup> kPa<sup>-1</sup> | 72 mm kPa<sup>-1</sup> |
| ETc · ETo<sup>-1</sup> | 0.30 | 12.6 | 0.0005 | 11.0 | 51.7 t ha<sup>-1</sup> | 0.265 |
| Rain (mm) | 0.62 | 76.45 | 0.0001 | 13.3 | 0.117 t ha<sup>-1</sup> mm<sup>-1</sup> | 74 mm |
| Rain · VPD<sup>-1</sup> (mm kPa<sup>-1</sup>) | 0.59 | 64.87 | 0.0001 | 13.3 | 0.26 t ha<sup>-1</sup> mm<sup>-1</sup> kPa<sup>-1</sup> | 22 mm kPa<sup>-1</sup> |
| Rain · ETo<sup>-1</sup> | 0.56 | 56.01 | 0.0001 | 13.3 | 88.1 t ha<sup>-1</sup> | 0.087 |

**Fig 4.**
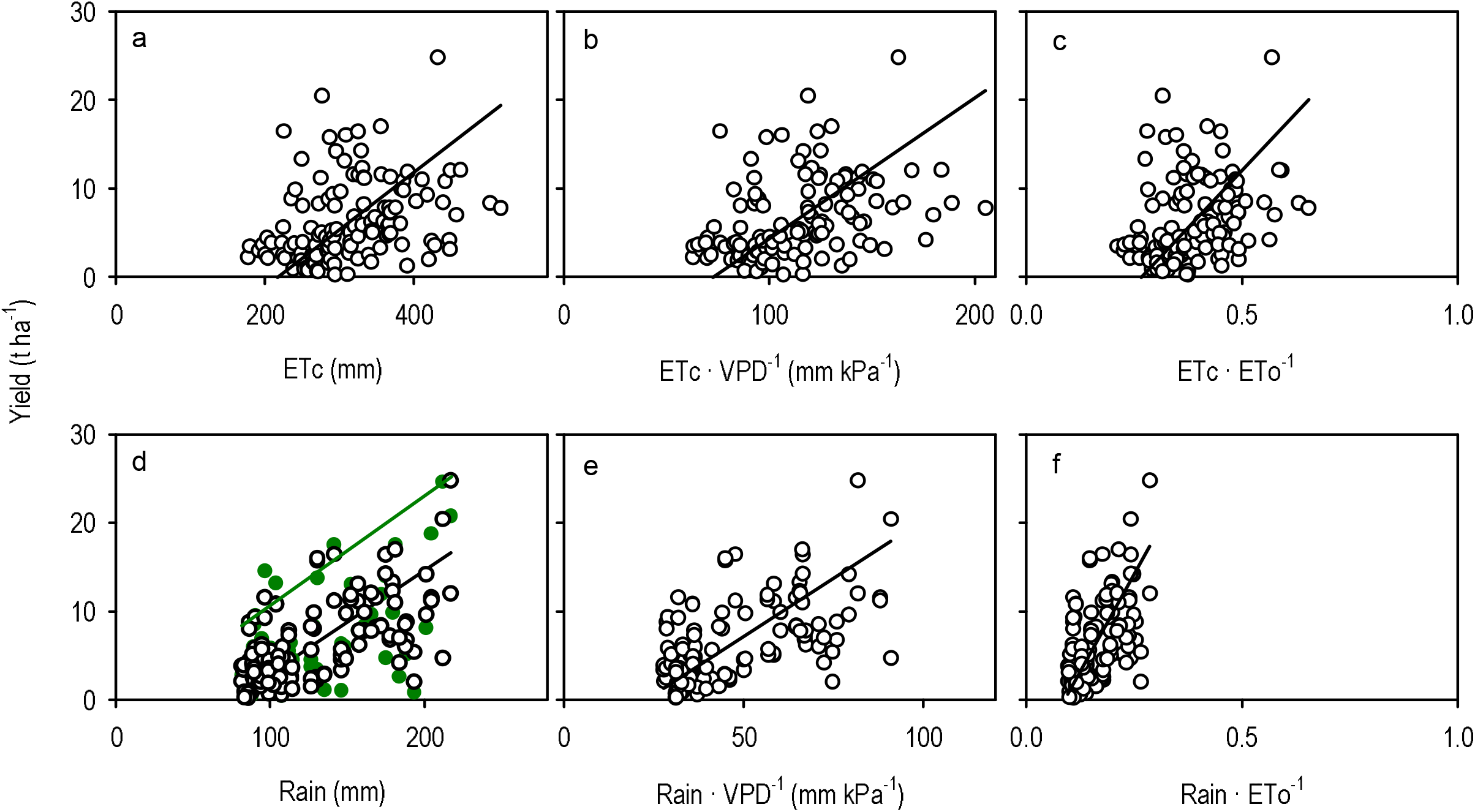
Relationship between vine yield and seasonal (a) crop evapotranspiration, ETc; (b) ETc corrected by vapour pressure deficit; (c) ETc corrected by reference evapotranspiration, ETo; (d) rainfall; (e) rainfall corrected by vapour pressure deficit; (f) rainfall corrected by ETo. Black lines are reduced maximum axis regressions fitted to a common data set (open black circles), with statistics summarised in Table 2. In (d), green symbols are additional data, and the green line is the 95^th^ percentile boundary derived from all data: *y = -1*.*92 + 0*.*124 x*.

We fitted a boundary function relating yield and seasonal rainfall for benchmarking vine yield (Fig. 4d, green line). Although seasonal rainfall is only one component of the total water input, we used rainfall instead of ETc for four reasons. First, seasonal rainfall returned the strongest correlation with yield (Table 2). Second, the spatial variation of seasonal rainfall in the Barossa regions correlates with the spatial variation in annual rainfall (Bramley and Ouzman 2021). Third, there were more data available to fit a boundary function with rainfall than for ETc; the association between yield and seasonal rainfall for the larger data set returned r = 0.59, F_1,188_ = 102.14, p < 0.0001, s > 13.3. Fourth, rainfall is more readily available than ETc for industry applications.

### Yield gap

The average yield gap was twice as large in 2021, the highest-yielding season, compared with 2019 and 2020 (Fig 5a). The average yield gap varied from 9.9 t ha^-1^ in the Central Grounds to 16.8 t ha^-1^ in the Southern Grounds (Fig. 5b). Part of this variation was related to elevation, with yield gap declining at 0.038 ± 0.01 t ha^-1^ m^-1^ in the range from 186 to 326 m.a.s.l. (Fig. 5c); the Eden Valley departed from this trend, with a yield gap that was higher than expected from its elevation (solid symbols in Fig. 5c).

**Fig 5.**
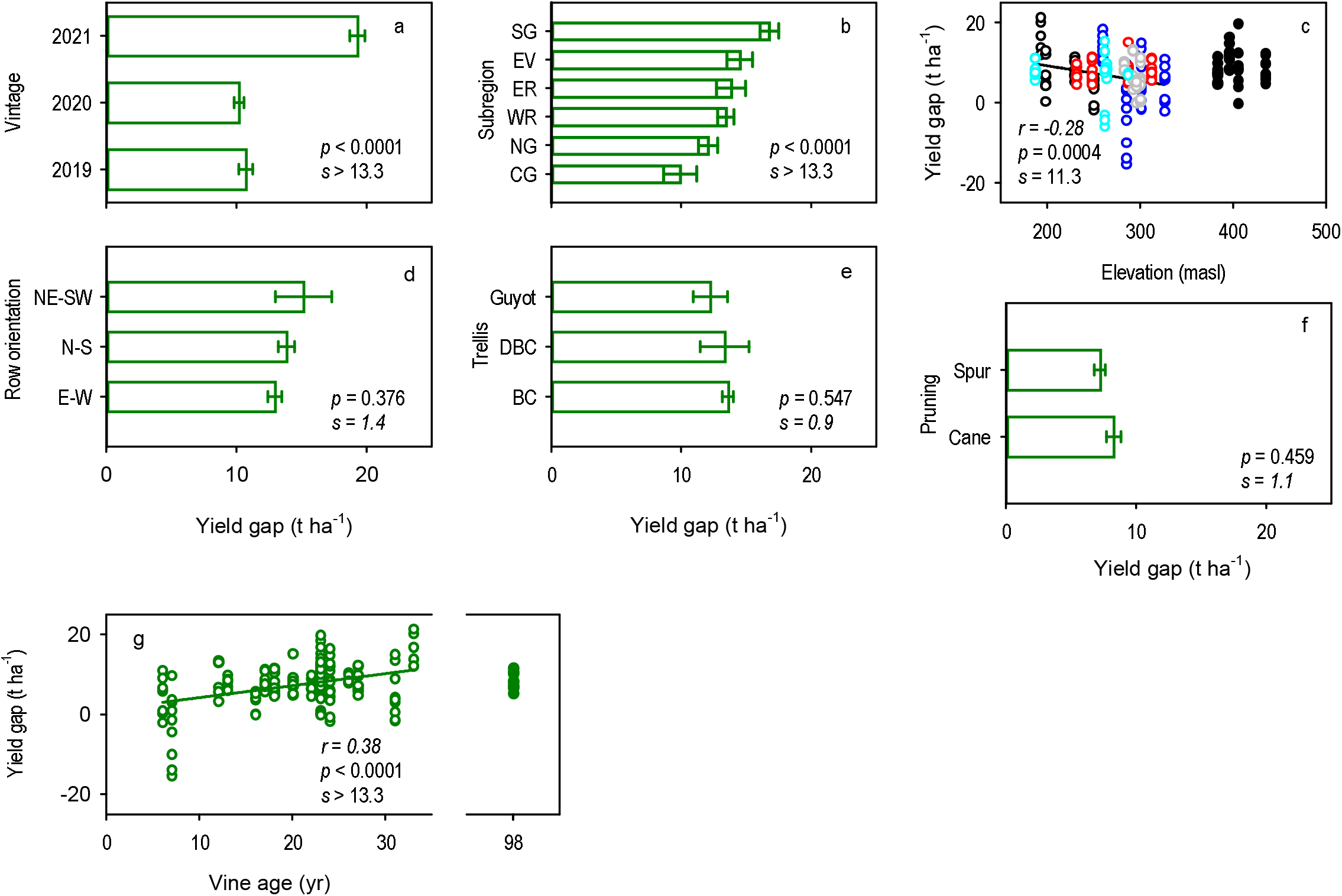
Variation in the yield gap of Shiraz in the Barossa zone (GI) with (a) vintage, (b) subregion, (c) elevation, (d) row orientation, (e) trellising system, (f) pruning method, and (g) vine age. In (a,b, d, e, f), error bars are two standard errors of the mean, and *p* and *s* from ANOVA. Subregions are Southern Grounds, shown as SG in (b) and open black symbols in (c), Eastern Edge (EE, blue symbols), Western Ridge (WR, red symbols), Northern Grounds (NG, gray), Central Grounds (CG, cyan), and Eden Valley (EV, closed black symbols). In (c), the line is the least squares regression fitted to all subregions but Eden Valley. In(f), the line is the least squares regression fitted to age between 6 and 33 years (open symbols); solid symbols (not included in the regression) are the yield gap of reworked vines on 98-year old rootstock.

The yield gap did not vary with trellising system, pruning method, or row orientation (Fig. 5def). The yield gap increased slightly with distance between vines at 3.0 ± 1.5 t ha^-1^ m^-1^ (r = 0.14, *p* = 0.051, s = 4.3), and did not vary with distance between rows (*p* = 0.208, s = 2.3) or rectangularity (*p* = 0.251, s = 2.0). The yield gap increased linearly with vine age between 6 and 33 yr at a rate of 0.3 ± 0.06 t ha^-1^ yr^-1^ (Fig. 5g). The yield gap in 98-year-old Shiraz that had recently been reworked averaged 8.3 ± 0.70 t ha^-1^ (solid symbols in Fig. 5g); this compares with a projected gap from the fitted regression of 31 t ha^-1^.

The yield gap correlated with all four yield components; the strength of the correlation ranked bunch weight ≈ berries per bunch > bunch number > berry weight (Fig. 6a-d). The minimum to close the yield gap was 185,000 bunches ha^-1^, 105 g bunch^-1^, 108 berries bunch^-1^ and 1.1 g berry^-1^ (arrow heads in Fig. 6a-d). The yield gap declined with increasing pruning weight, and the minimum pruning weight to close the gap was 3.2 t ha^-1^.

**Fig 6.**
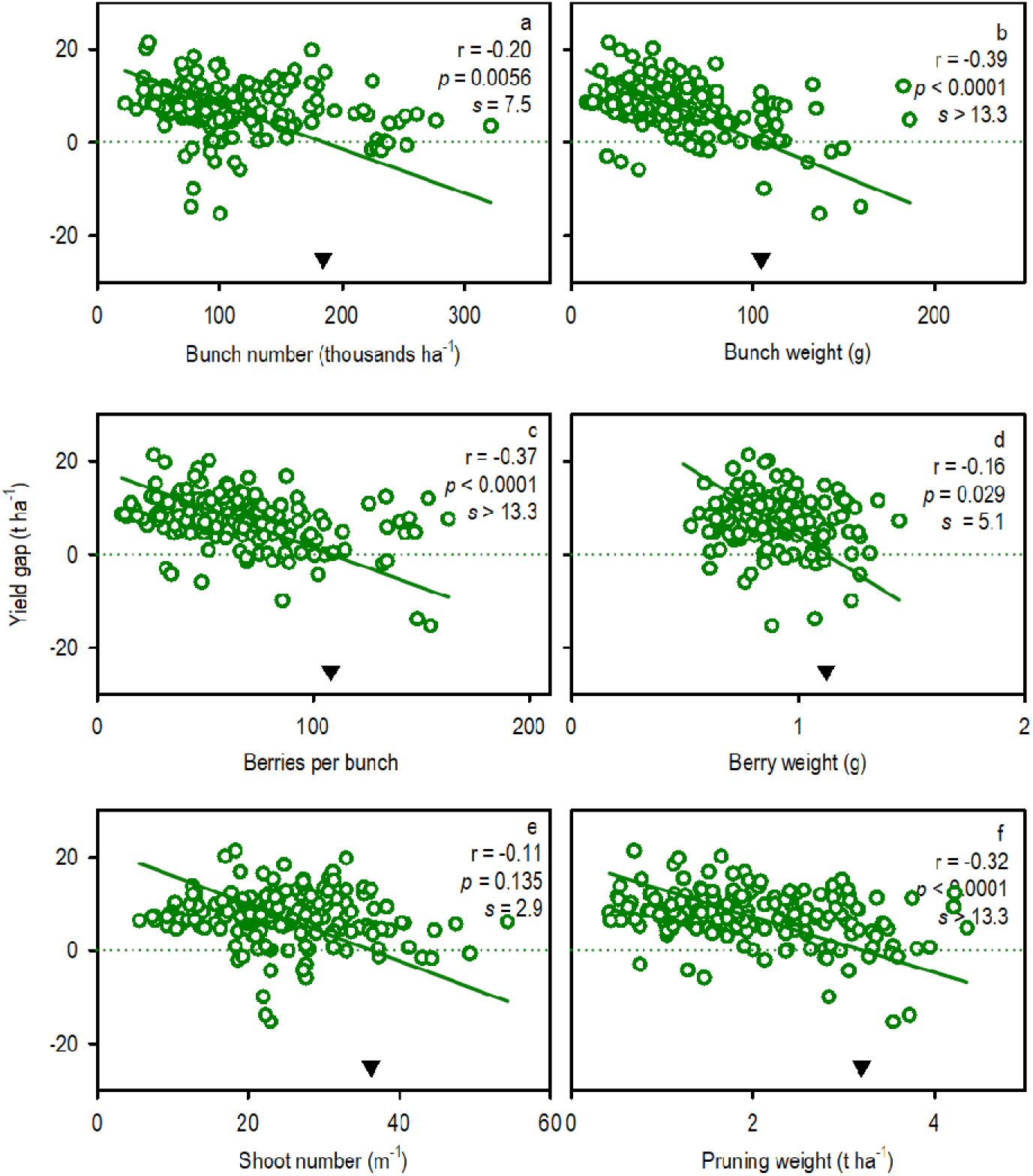
Relations between yield gap of Shiraz in the Barossa zone (GI) and (a) bunch number, (b) bunch weight, (c) berries per bunch, (d) berry weight, (e) shoot number, and (f) pruning weight. Lines are reduced maximum axis regressions. Arrowheads show the trait value for yield gap = 0.

Pearson’s correlation coefficients were calculated between the yield gap and weather factors in four periods, from late August to budburst, budburst to flowering, flowering to veraison, and veraison to maturity (Fig. 7, and correlation matrix for all factors Supplementary Table 1). The association between yield gap and both maximum and minimum temperature shifted from positive before budburst to negative after flowering. The association between yield gap and radiation was negative from budburst to flowering and from veraison to maturity, and positive in the intervening period from flowering to veraison. Larger yield gaps were associated with smaller VPD between budburst and maturity, and the association was stronger at earlier stages. The association between yield gap and rainfall was negative before budburst and between flowering and veraison, and positive between budburst and flowering. The larger yield gap with higher rainfall between budburst and flowering was partially associated with the strong negative correlation between rainfall and VPD (Supplementary Table 1). The yield gap increased with increasing vine water stress quantified with carbon isotope composition of must (Fig. 2c).

**Fig 7.**
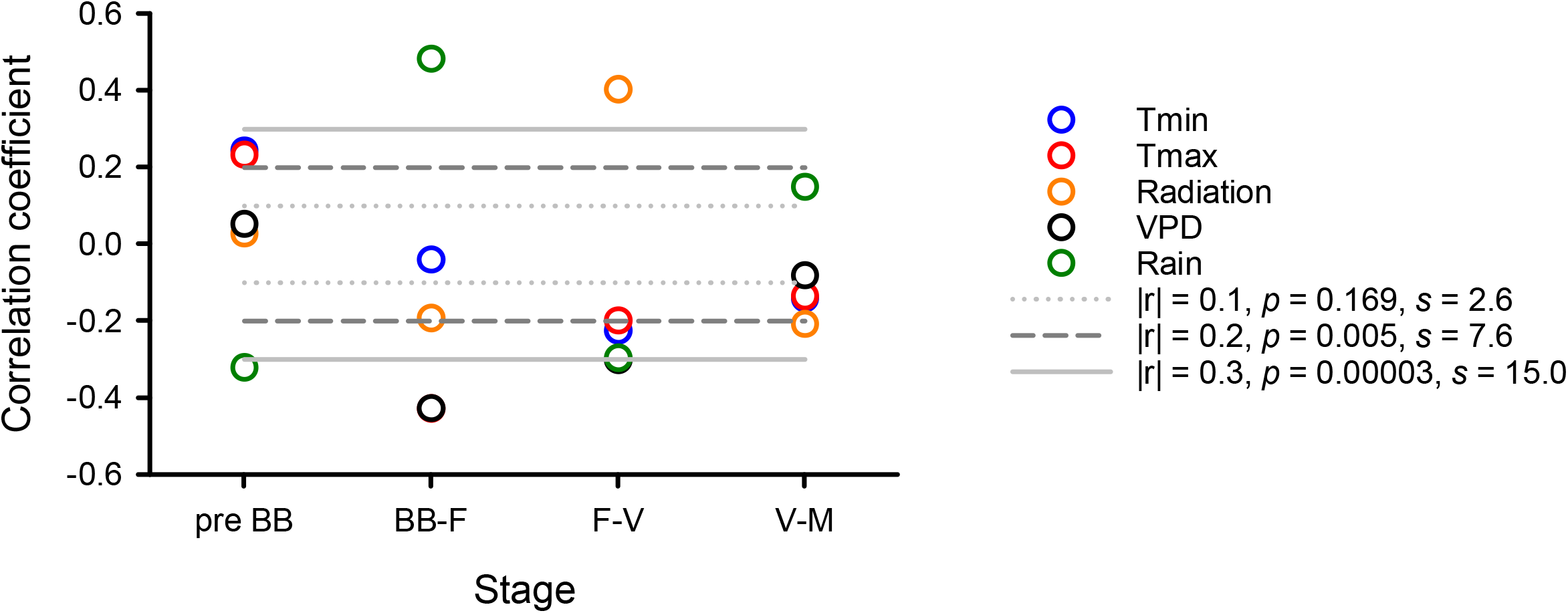
Correlation coefficient for the association between yield gap and meteorological variables from 29 August to budburst to budburst (pre-BB), from budburst to flowering (BB-F), from flowering to veraison (F-V), and from veraison to maturity (V-M).

## Discussion

Climate change is placing increased stress on water resources and supports a revisions of water management strategies and how they relate to vineyard productivity. This will help account for direct (i.e., shifts in amount and seasonality of rainfall, reduced availability of water for irrigation, and higher temperature and vapour pressure deficit), and indirect effects of climate change including the use of irrigation to manage heat stress. In this context, we benchmarked the relation between yield and water use, calculated yield gaps, and explored their underlying environmental and management causes.

The reduced-maximum axis regression between yield and ETc returned a slope of 0.065 t ha^-1^ mm^-1^ and x-intercept of 216 mm (Table 2). This compares with slopes from 0.061 to 0.123 t ha^-1^ mm^-1^ and x-intercepts from 136 to 175 mm in a sample of vineyards in Spain, the US and China (Table 3). Parameters from least squares regressions, the default in most software packages, are included in Table 3 to highlight the flatter slopes and lower x-intercepts from this approach that assumes error in x is negligible in relation to error in y (Ludbrook 2012, Niklas 1994); the assumption is unjustified because error in ETc is not negligible. Indeed, yield correlated more strongly with seasonal rainfall than with ETc (Fig. 4; Table 2). This was related to the uncertainty in the quantification of both soil water content and irrigation used in the calculation of ETc (Vinod et al. 2022 pre-print). We thus favoured a rainfall-based benchmark for water-limited yield potential and yield gap analysis. This model has several assumptions, including a single x-intercept representing soil evaporation. The model developed by Rockström (2003) explicitly accounts for the reduction in soil evaporation associated with larger canopies and higher yield (eq. 2). Our data conformed to this model, but statistical and biophysical criteria supported the conclusion that it is unsuitable for our analysis: the relationship ETc:yield vs yield is spurious (Brett 2004), and inconsistent with the small variation in Es:ET with increasing yield (Fig. 3).

**Table 3.**
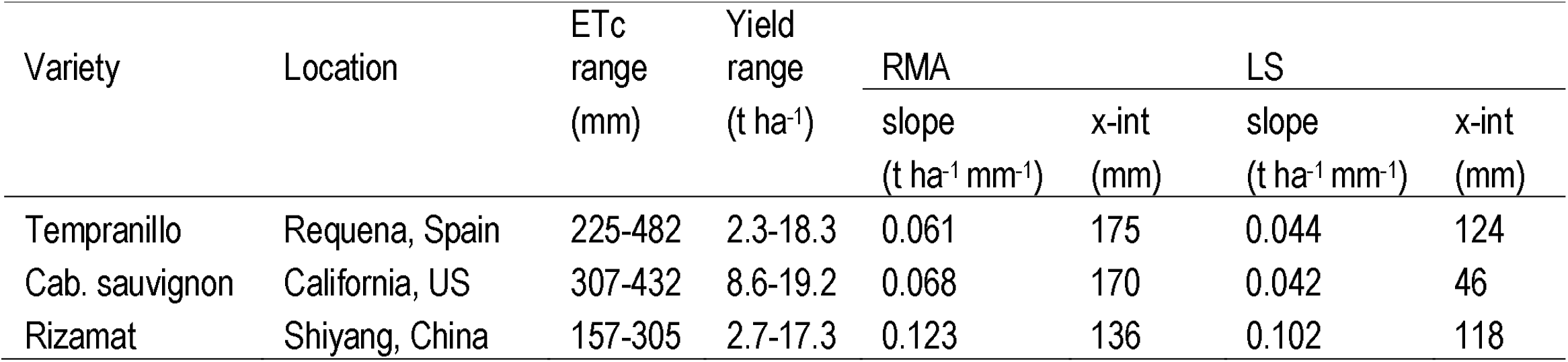
Range of crop evapotranspiration (ETc) and yield, and slopes and x-intercepts from regressions between yield and ETc in a sample of vineyards from the literature. Regression methods are reduced-maximum axis (RMA, Model II) and least squares (LS, Model I) for comparison. Calculated from data reported by Intrigliolo and Castel (2008) (Spain), Williams (2010) (US) and Du et al. (2008) (China).

In contrast to annual crops where canopy size is a large source of variation in energy partitioning, the wide-row structure dampens the variation in Es:ETc of vineyards, particularly in low-rainfall environments and under drip irrigation that wets a limited soil area (Jiao, et al. 2018, Yunusa, et al. 1997, Yunusa, et al. 2004, Zhang, et al. 2010). In a furrow-irrigated Sultana vineyard in the arid Victorian Mallee, Australia, Es:ETc was 0.46-0.51 for a yield range between 7 and 22 t ha^-1^ (Yunusa, et al. 1997). In a drip-irrigated Sultana vineyard in the same region, with a yield range from 6 to 20 t ha^-1^, the RMA rate of decline in Es:ETc with increasing yield was 0.011 ± 0.003 (t ha^-1^)^-1^ (Yunusa, et al. 1997) compared with the rate of 0.018 ± 0.002 (t ha^-1^)^-1^ in our data set (Fig. 3b). These two studies (Yunusa, et al. 1997, Yunusa, et al. 1997) are thus consistent with both the flat relation between Es:ETc and yield in vineyards, and the lower Es:ETc at high yield under drip irrigation compared to furrow irrigation.

The spatial coherence of seasonal and annual rainfall in the Barossa regions (Bramley and Ouzman 2021) and the biophysical and viticultural meaningful yield gaps derived from a rainfall-based benchmark reinforce the robustness of our approach. The yield gap was larger in the season with highest yield and varied with location (Fig. 5ab). Elevation, a major factor in the clustering of locations in the Barossa regions (Bramley and Ouzman 2021), accounted for part of the variation with smaller gaps with increasing elevation (Fig. 5c). Historically, the name Barossa Valley was used to describe the area below 400 masl, and the name Eden Valley, rather than Barossa Ranges, has been favoured for higher elevations since the 1950s (Fuller and Walsh 1999). The Eden Valley locations departed from this trend, with larger yield gaps than expected from their elevation (Fig. 5c). This larger-than-expected yield gaps can be related management shallow soil and limited access to irrigation water (Bramley and Ouzman 2021, Department of Environment and Water 2022). Unlike the Barossa Valley region that relies largely on irrigation water from the River Murray, the Eden Valley region is heavily reliant on native sources, i.e., groundwater and surface water (Department of Environment and Water 2022).

The yield gap did not vary with row orientation, trellis system or pruning method (Fig. 5). Water use of vertically trained, potted vines was approximately 18% lower in east-west orientated vines compared to north-south vines in the northern hemisphere (Buesa, et al. 2020). Row orientation did not affect carbon assimilation; hence, the water use efficiency (based on both carbon assimilation and yield) was higher for the east-west vines was higher than the vines orientated north-south (Buesa et al 2020). Modelled intercepted radiation is higher in north-south than east-west orientated rows (Palmer 1989). However, the impact of row orientation is complicated by its interactions with and between canopy configuration, time of year, and row spacing (Jackson and Palmer 1972) as well as vine water status, crop load and the other practices discussed above. Shiraz vines in the Barossa and Eden Valley regions are traditionally managed using spur pruning and a single cordon, with some vineyards maintaining a second cordon approximately 0.4 m above the first (Dry, et al. 2005). More recently, occasional vineyards have been changed to Guyot, locally described as rod or cane pruning, often to improve trunk disease management (Henderson, et al. 2021). Multiple cordons increase the number of buds retained on the vine and canes favour bud fertility, hence both systems have the potential to increase bunch number and yield relative to the single cordon spur pruned system (Jackson 2001).

The yield gap increased linearly with vine age, and this can be related to at least four factors: time to peak production, technological change, disease progression and vine management. Crop evapotranspiration and yield increase with vine age from establishment until the canopy and root system reach their full capacity to capture radiation and water, around 6 to 10 years after planting depending on management, variety and growing conditions (Evans, et al. 1993, Munkvold, et al. 1994, Sadras and Schultz 2012). Hence, age-related variation in vine capacity to capture water and radiation was a minor factor for the range from 6-to 33-year-old vines in which yield gap related to vine age. Eutypa dieback, caused by the fungus *Eutypa lata*, is a severe, widespread trunk disease of grapevines (Koussa, et al. 2006, Munkvold, et al. 1994, Sosnowski, et al. 2008). The fungus grows slowly in the plant and the onset of foliar symptoms occurs 3-8 years after infection. In south-eastern Australian vineyards, the incidence of trunk disease increased with vine age (Sosnowski, et al. 2016). The yield of Chenin blanc vineyards in California varied non-linearly with age, increasing up to a peak over 20 t ha^-1^ in 12-year-old vines, and declined dramatically after this peak in association with the period of rapid increase in Eutypa dieback (Munkvold, et al. 1994). The putative effects of disease can be partially confounded with management; arm wrapping into the cordon wire is a practice commonly used during the establishment of a new vineyard, which may bring a de-vigorating effect and potential drop in production as cordon strangulate over the years (O’Brien, et al. 2021). Irrespective of the causes, the costs of dealing with older, less productive vines (vine redevelopment, new planting, and delay of productivity of young vines) need to be weighed against the rate of increase in yield gap of 0.3 ± 0.06 t ha^-1^ yr^-1^ (Fig. 5g).

Plant water deficit was neutral or slightly improved the tolerance of grapevine to infection and colonisation by E. lata (Sosnowski, et al. 2021). Once established in the plant, Eutypa dieback fungi invade bark tissues and xylem, resulting in the death of a portion of vascular cambium whereby infected structures are no longer able to produce newly functional xylem and phloem, and cankers develop (Pouzoulet, et al. 2014). Eutypa dieback reduced the water content and disrupted the dynamics of free abscisic acid (ABA) and its glucose esters (ABA-GE) in diseased organs of Cabernet sauvignon (Koussa, et al. 2006). Consistent with the disruption in the vascular system and altered water relations, the yield gap increased with water stress, as quantified with carbon isotope composition (Fig. 2c). Other factors that contribute to water stress and yield gap include soil plant available water, primarily related to soil depth (Bramley and Ouzman 2021, Coipel, et al. 2006).

Yield correlated with all four yield components, particularly with bunch number (Table 1); bunch number is usually the largest source of variation in grapevine yield (Dunn, et al. 2004). The yield gap correlated with all four yield components, particularly bunch weight and berries per bunch, highlighting the incidence of berry set. The causes for the shift from bunch number as the main source of variation in yield to bunch size as the main source of variation in yield gap are unclear. Irrespective of the causes, the minimum bunch size (105 g bunch^-1^, 108 berries bunch^-1^) to close yield gaps maybe used as a rule-of-thumb to benchmark vineyard performance, but requires independent testing (Fig. 6). The yield gap was smaller with higher rainfall before budburst, putatively favouring early-season vegetative growth and allocation to reproduction, and with higher rainfall between flowering and veraison, putatively favouring fruit set and early berry growth (Friend, et al. 2009, Ristic and Iland 2005). Consistent with our correlative findings, experimental evidence led to the conclusion that winter irrigation is required to maintain yield in the context of drier winters with climate change (Bonada, et al. 2020). The gap was larger with high rainfall, and associated low radiation, between budburst and flowering (Fig. 7). For Godello in the Ribero Designation of Origin, the production of pollen spanned 2-3 weeks in May and June, and yield correlated negatively with rainfall in May (González-Fernández, et al. 2020). Three successive rainy days in late May (53 mm), just after the seasonal pollen peak, promoted a fast decrease in the airborne pollen concentration (González-Fernández, et al. 2020). The association between yield gap and temperature shifted from positive early in the season, to negative at later stages, particularly between flowering and veraison. This agrees with empirical evidence that indicates differential sensitivity to temperature between phenological stages. Consistent with the positive correlation between yield gap and temperature early in the season observed here, artificially elevated temperature two weeks before budburst decreased the number of flowers at a rate of –7.3 flowers per ºC for the lower inflorescence and – 5.5 flowers per ºC for the upper inflorescence, with no impact of temperature two weeks after budburst (Petrie and Clingeleffer 2005). Temperature sensitivity increases again during anthesis; the damage, however, depends on the threshold of temperature. Temperature over 35 °C at flowering reduced fruit set and the number of berries per bunch (Kliewer 1977), whereas temperature around 28 °C favour pollen germination and ovule fertilization (Staudt 1982). Therefore, we hypothesise that higher temperature from flowering to veraison during the mild seasons of this experiment may have contributed to reduce cold damage during flowering, contributing to increase fruit set and closing the yield gap.

## Acknowledgements

We thank Gaston Sepulveda, Annette James, and Han Chow for vineyard and laboratory work, Tony Hall and the Mawson Analytical Spectrometry Services (MASS) Facility at the University of Adelaide for the analysis of carbon isotopes, Wine Australia for funding project UA1602, and our project partners. Wine Australia invests in and manages research, development and extension on behalf of Australia’s grapegrowers and winemakers and the Australian Government. The South Australian Research and Development Institute is a member of the Wine Innovation Cluster in Adelaide.

